# Lipid droplets are a metabolic vulnerability in melanoma

**DOI:** 10.1101/2022.05.04.490656

**Authors:** Dianne Lumaquin, Emily Montal, Arianna Baggiolini, Yilun Ma, Charlotte LaPlante, Ting-Hsiang Huang, Shruthy Suresh, Lorenz Studer, Richard M. White

## Abstract

Melanoma exhibits numerous transcriptional cell states including neural crest-like cells as well as pigmented melanocytic cells. How these different cell states relate to distinct tumorigenic phenotypes remains unclear. Here, we use a zebrafish melanoma model to identify a transcriptional program linking the pigmented cell state to a dependence on lipid droplets, the specialized organelle responsible for lipid storage. Single-cell RNA-sequencing of these tumors show a concordance between genes regulating pigmentation and those involved in lipid and oxidative metabolism. This state is conserved in human melanoma specimens. This state demonstrates increased fatty acid uptake, an increased number of lipid droplets, and dependence upon oxidative metabolism. Genetic and pharmacologic suppression of lipid droplet production is sufficient to disrupt oxidative metabolism and slow melanoma growth in vivo. Because the pigmented cell state is linked to poor outcomes in patients, these data indicate a metabolic vulnerability in melanoma that depends on the lipid droplet organelle.

## Introduction

Phenotypic heterogeneity is influenced both by cell intrinsic factors and tumor microenvironmental (TME) interactions^1–4^. In melanoma, both bulk and single-cell RNA-sequencing (scRNA-seq) has revealed numerous transcriptional cell states linked to distinct phenotypes^1–5^. For example, early studies demonstrated two unique gene signatures associated with proliferative versus invasive cell states^6,7^. These signatures have become increasingly nuanced, with most human melanoma exhibiting multiple transcriptional states linked to phenotypes such as metastasis, drug resistance and immune evasion^2,3,8–10^. While these states are somewhat plastic, their maintenance across different patients suggests that each state may exhibit unique biological properties. One dimension of this phenotype spectrum is the degree of cellular differentiation. Melanocytes arise from the progressive differentiation of neural crest cells, to melanoblasts, and finally differentiated melanocytes^11^. scRNA-seq has revealed that most melanomas contain a mixture of undifferentiated neural crest-like cells as well as more mature, pigmented melanocytic cells^2,4^. The less mature, neural crest-like states have been implicated in melanoma initiation and metastasis and have been well characterized^12,13^. In contrast, fewer studies have focused on the role of the pigmented, mature melanocytic cell state. Prior to terminal differentiation, these pigmented melanocytic cells remain highly proliferative, suggesting they may play a key role in tumor progression^9,14,15^. This state has translational importance, since clinical data has correlated differentiated melanocytic identity with worse clinical outcomes^16–19^. Acquisition of a differentiated melanocytic cell identity has been associated with metastatic seeding and evasion to targeted therapy in patients, highlighting the need to uncover vulnerabilities in this cell state^19,20^.

Dissecting the mechanisms driving this state requires tractable genetic models that recapitulate human melanoma phenotypic heterogeneity while preserving an immunocompetent TME. In this study, we use a zebrafish model of BRAF-driven melanoma coupled with analysis of human samples to uncover a metabolic dependency of the pigmented melanocytic state that is linked to lipid droplets.

## Results

### Melanocytic cell state upregulates oxidative metabolism in melanoma

We used our recently developed technique for generating melanoma in zebrafish, TEAZ (Transgene Electroporation of Adult Zebrafish)^21^, to investigate phenotypic heterogeneity in a model of *BRAF*^*V600E*^ *p53^-/-^ PTEN^ko^* melanoma (Fig. 1a). In this model, fish develop melanomas in fully immunocompetent animals at the site of electroporation in a median of 5 to 8 weeks. We performed scRNA-seq from n=6 zebrafish tumors for a total of 3968 cells composed of melanoma and TME cells (Fig. 1b). After quality control and cluster annotation^22^, we identified five melanoma clusters representing distinct transcriptional cell states (Fig. 1b,c and Supplementary Fig. 1 a-e). Using differential gene expression and Gene Set Enrichment Analysis (GSEA), we identified the five melanoma clusters to represent transcriptional states of stressed, proliferative, invasive, inflammatory, and melanocytic identity (Fig. 1c, Supplementary Fig. 2 a-d and Supplementary Data 1). The stressed cluster in our data set significantly enriched for genes like *gadd45ga*, *ddit3*, *fosb*, and *junba*, consistent with the stressed cellular state in previous zebrafish and human melanoma scRNA-seq profiling^2,8^ (Fig. 1c and Supplementary Fig. 2a). Through GSEA, we found that the inflammatory cluster enriched for immune related processes and the proliferative cluster strongly enriched for cell division processes (Fig. 1c and Supplementary Fig. 2b-c). Similarly, the invasive cluster enriched for genes like *krt18a.1* and *pdgfbb* expressed in previously characterized human melanoma invasive cell states^7^, as well as cell migration processes (Fig 1c and Supplementary Fig. 2d),

**Fig. 1:**
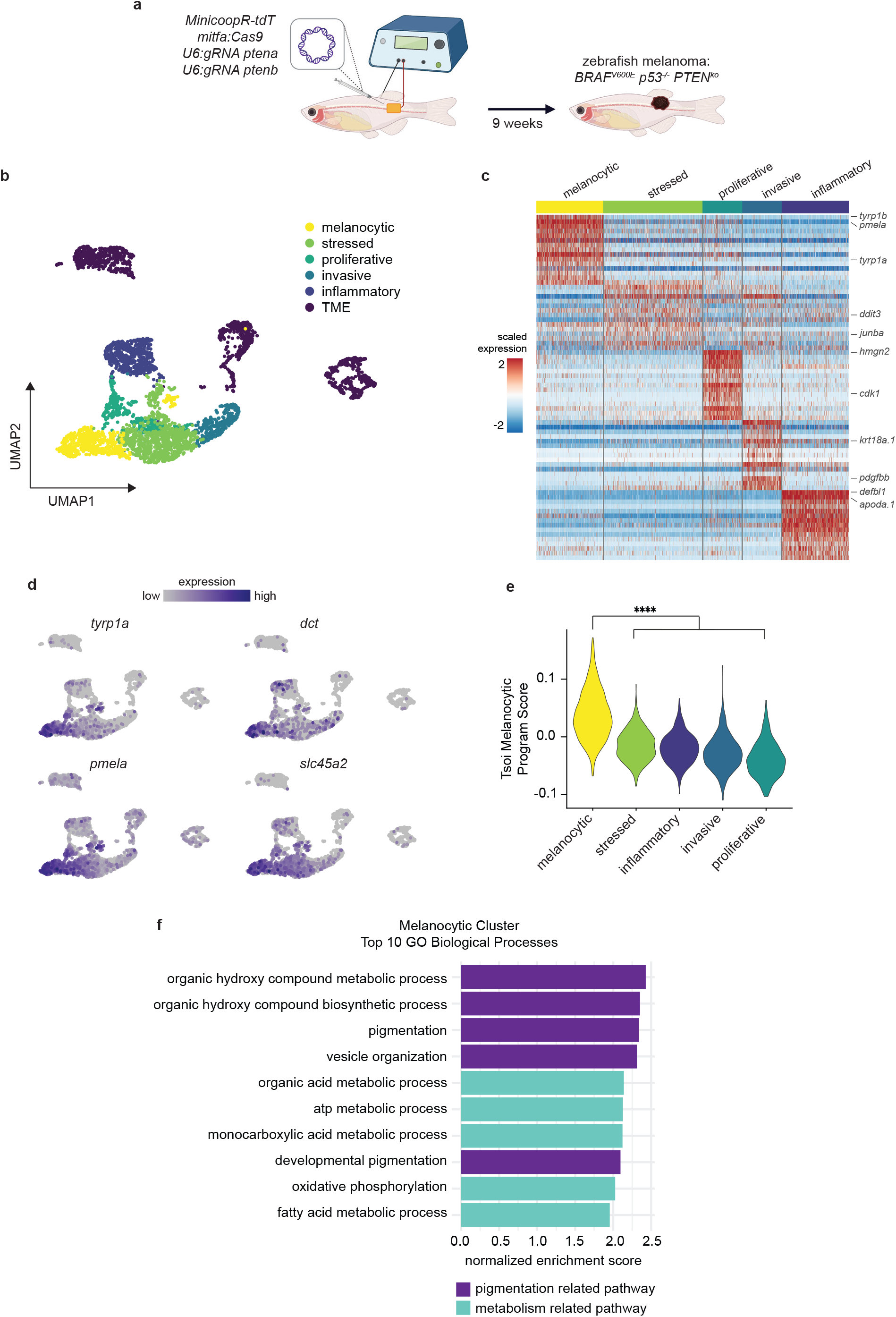
Zebrafish melanoma displays distinct transcriptional states where melanocytic cell state upregulates oxidative metabolic pathways. a, Schematic of zebrafish TEAZ. Transgenic *casper*;*mitfa*:*BRAF^V600E^*;*p53^-/-^*zebrafish were injected with tumor initiating plasmids and electroporated to generate *BRAF^V600E^ p53^-/-^ PTEN^ko^* melanomas. b, UMAP dimensionality reduction plot of melanoma and TME cells. Cell assignments are labeled and colored. c, Heatmap of top 15 differentially expressed genes in each melanoma transcriptional cell state. Select genes labeled. d, UMAP feature plots showing scaled gene expression of pigmentation genes (*tyrp1a*, *dct*, *pmela*, *slc45a2*). e, Tsoi Melanocytic Program^4^ module scores for cells in each melanoma cell state. Adjusted p-values calculated using Wilcoxon rank-sum test with Holm correction. f, Top 10 enriched GO Biological Processes in melanocytic cluster with Benjamini-Hochberg adjusted p-values<0.05. Pathways are color coded based on pigmentation or metabolism related pathways.

Differentiated melanocytes are marked by expression of pigmentation genes involved in melanin production^11,23^. The differentiated melanocytic cluster in our dataset showed enrichment for pigmentation genes expressed by differentiated melanocytes such as *tyrp1a*, *dct*, *pmela*, and *slc45a2* (Fig. 1c-d). To compare our zebrafish data to human data, we used module scoring to compare gene enrichment to the human melanocytic signature from Tsoi et al.^4^ and found the zebrafish melanocytic cluster to enrich for this differentiated melanocytic gene program (Fig. 1e). When we performed GSEA on the melanocytic cluster, we found an enrichment for pigmentation related pathways among the top 10 GO Biological Processes as expected (Fig. 1f). Aside from this, we also noted that this cluster enriched for oxidative phosphorylation and fatty acid metabolism. To determine if this was unique to the zebrafish, we also analyzed the human melanoma brain and leptomeningeal metastases scRNA-seq dataset from Smalley et al.^24^ (Supplementary Fig. 3a). We found that the cluster most significantly enriched for the Tsoi melanocytic gene program^4^ also displayed enrichment for oxidative metabolism transcriptional programs (Supplementary Fig. 3b-c). Altogether, these data suggest that acquisition of the melanocytic cell state correlates with a gene signature of oxidative metabolism.

To functionally test whether cells become more oxidative as they adopt a melanocytic cell state, we used a human pluripotent stem cell system. Either embryonic stem (ES) or induced pluripotent stem (iPS) cells can be differentiated into neural crest cells, melanoblasts or mature melanocytes^11,13^. Similar to our zebrafish model, we introduced the *BRAF^V600E^* oncogene and inactivated *PTEN* in the human pluripotent stem cells (Supplementary Fig. 4a-b). We then used the Seahorse Mito Stress Test to measure cellular OCR (oxidative consumption rate) and ECAR (extracellular acidification rate) in melanoblasts versus melanocytes (Fig. 2a,b). These are markers for oxidative and glycolytic metabolism, respectively. We observed a robust increase in the basal OCR and OCR/ECAR ratio in the melanocytes compared to the melanoblasts, suggesting elevated oxidative metabolism in these more mature cells (Fig. 2b). Similarly, we pharmacologically induced the melanocytic cell state in human A375 cells through increasing cAMP signaling via IBMX and Forskolin^25^ (Fig. 2d). Treatment with IBMX and Forskolin resulted in upregulation of pigmentation genes, *dct* and *pmel*, and concurrent increases in the basal OCR and OCR/ECAR ratio (Fig. 2e). Collectively, these data provide evidence for oxidative metabolic rewiring in melanoma cells adopting a pigmented, melanocytic cell state.

**Fig. 2:**
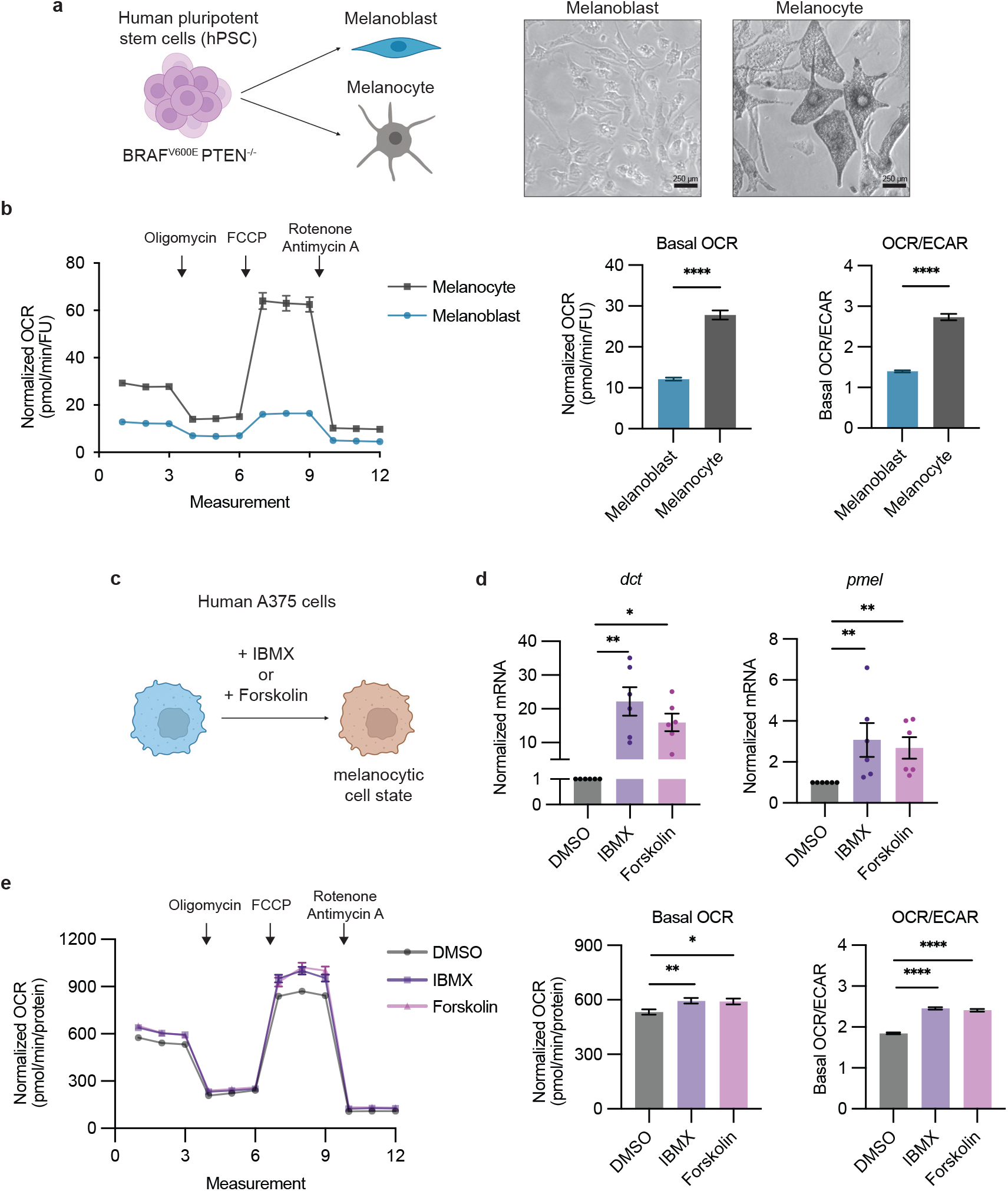
Melanocytic cell state leads to oxidative metabolic rewiring. a, Schematic of hPSC differentiation to melanoblasts and melanocytes. Representative images for cell morphology of melanoblasts and melanocytes. b, Normalized OCR measurements for melanoblasts and melanocytes. Basal OCR and OCR/ECAR values derived from measurement 3 (mean ± SEM, n = 3 biologically independent experiments). Statistics via two-tailed t-test with Welch correction. c, Schematic of inducing melanocytic cell state in human A375 cells via IBMX or Forskolin for 72 hours. d, qRT-PCR for melanocytic genes *dct* and *pmel* in human A375 cells (mean ± SEM, n = 3 biologically independent experiments). Statistics via Kruskal Wallis with Dunn’s multiple comparisons test. e, Normalized OCR measurements for human A375 cells treated with DMSO, IBMX or Forskolin. Basal OCR and OCR/ECAR values derived from measurement 3 (mean ± SEM, n = 3 biologically independent experiments). Statistics via One-way ANOVA with Dunnett’s multiple comparisons test. * p<0.05, ** p<0.01, **** p<0.0001.

### Fatty acids are fuel for oxidative metabolism and stored in lipid droplets

Oxidative metabolism can be fueled by various substrates including lipids, glucose and glutamine^26^. Since we observed an enrichment for fatty acid pathways in the zebrafish melanocytic cluster, we next asked whether fatty acids were increasingly utilized as substrates for β-oxidation in the melanocytic cell state (Fig. 1f). To test this, we performed the Seahorse Fatty Acid Oxidation (FAO) assay in human A375 melanoma cells +/- IBMX or Forskolin to induce melanocytic differentiation in the presence of oleic acid (Fig. 3a and Supplementary Fig. 5a-b). We observed higher OCR with oleic acid supplementation in the melanocytic cells (compared to control A375 cells), which is lost upon inhibition of FAO using etomoxir (Fig. 3a and Supplementary Fig. 5c). This data suggested an increase in FAO in cells adopting a melanocytic cell state. FAO can be fueled by either de novo synthesis or uptake from extracellular sources, and previous studies have shown that melanoma cells can upregulate exogenous fatty acid uptake through fatty acid transporter proteins^27,28^. To assess this in the melanocytic state, we added fluorescently labeled fatty acids to the media and measured fluorescence intensity as an indicator of fatty acid uptake into the cells^27,28^. This revealed that the more melanocytic cells increase fatty acid uptake compared to control A375 cells (Fig. 3b).

**Fig. 3:**
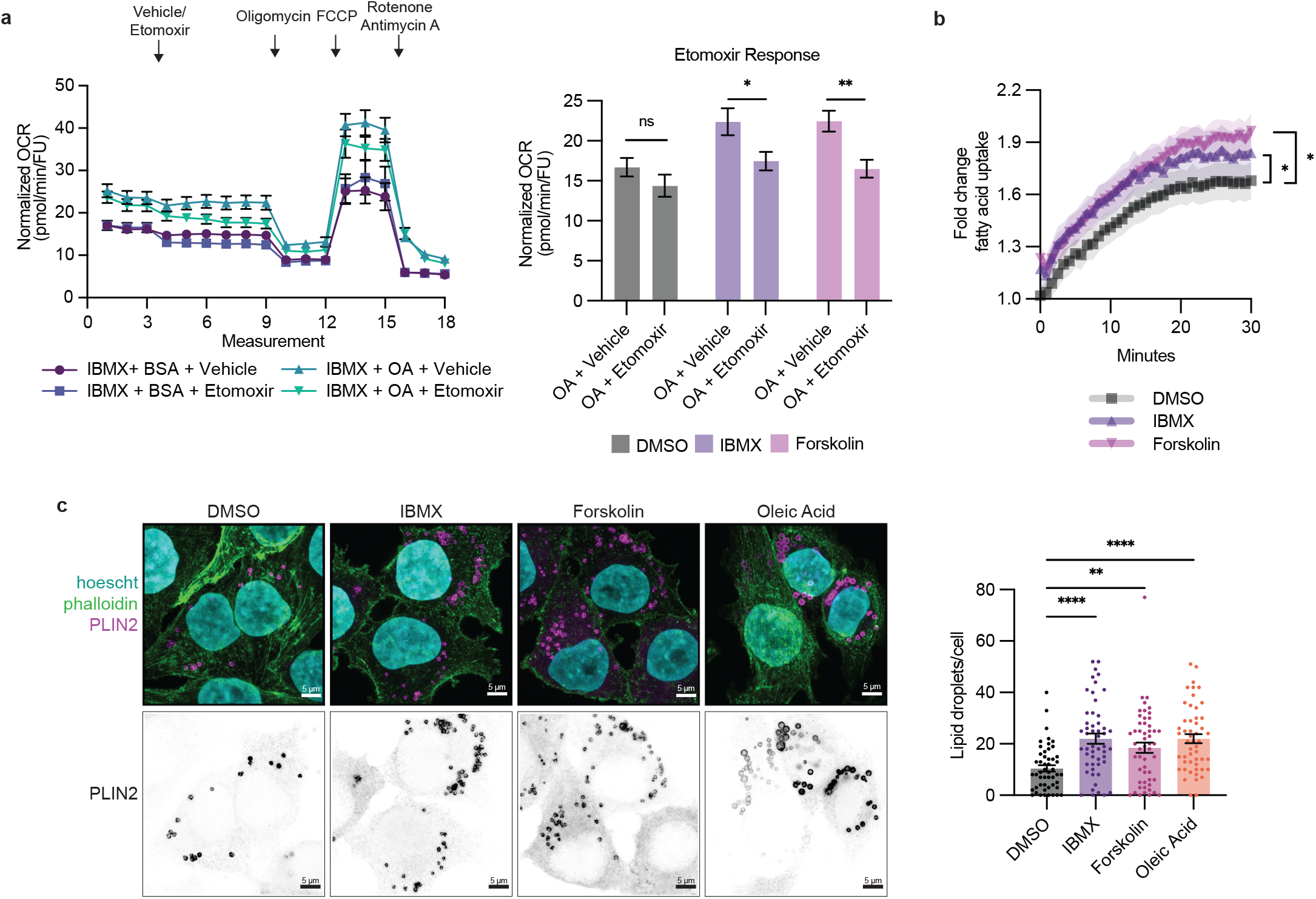
Fatty acids are utilized for oxidative metabolism and stored in lipid droplets. a, Representative normalized OCR measurements for FAO stress test in IBMX treated human A375 cells. Etomoxir response derived from normalized OCR values from measurement 9 (mean ± SEM, n = 3 biologically independent experiments). Statistics via two-tailed t-test with Holm-Sidak correction. b, Fold change in fatty acid uptake in drug treated human A375 cells (mean ± SEM, n = 3 biologically independent experiments). Statistics via area under the curve two-tailed t-test with Holm-Sidak correction. c, Representative fluorescent images of PLIN2+ lipid droplets in drug treated human A375 cells and corresponding quantification of PLIN2+ lipid droplets per cell (mean ± SEM, n = 3 biologically independent experiments). Statistics via Kruskal Wallis with Dunn’s multiple comparisons test. * p<0.05, ** p<0.01, *** p<0.001, **** p<0.0001.

While fatty acids can undergo metabolism through β-oxidation, excess levels of free fatty acids are toxic to cells and can limit proliferation, a phenomenon called lipotoxicity^29^. A major mechanism for avoiding such toxicity and maintaining proliferation is to package fatty acids as triacylglycerols in lipid storage organelles called lipid droplets^30^. Lipid droplets regulate lipid availability and shuttle fatty acids to the mitochondria for β-oxidation^31,32^. To test whether the more melanocytic cells utilized this mechanism of lipid storage, we stained the cells with an antibody against PLIN2, a major lipid droplet protein that we and others have previously shown marks this organelle in melanoma cells^33–35^. We quantified the number of lipid droplets per cell after treatment with oleic acid (as a positive control) or after treatment with IBMX or Forskolin to induce the melanocytic state. This showed a significant increase in the number of lipid droplets in the melanocytic state compared to control cells (Fig. 3c). Taken together, these data suggest that melanoma cells in the pigmented, melanocytic state undergo metabolic rewiring to increase fatty acid uptake and β-oxidation, while at the same time increasing lipid droplet numbers.

### Loss of lipid droplets suppresses melanoma progression and disrupts metabolic homeostasis

Lipid droplet accumulation has been associated with increased melanoma cell proliferation and invasion leading to poor clinical outcomes^27,35^. Given the evidence linking the melanocytic cell state and lipid droplet accumulation to worse clinical outcomes, we next asked if disrupting lipid droplet formation would affect melanoma progression. To test this, we focused on DGAT1, a critical enzyme in triacylglycerol synthesis that is well known as a target to inhibit lipid droplet biogenesis^32,36^. More recently, DGAT1 has been linked to reducing oxidative stress and lipotoxicity in glioblastoma and melanoma^37,38^. To determine whether loss of DGAT1 would perturb lipid droplet formation in melanoma cells, we used a DGAT1 inhibitor and CRISPR/Cas9 to knockout *DGAT1* in our zebrafish melanoma lipid droplet reporter^33^ (Fig. 4a and Supplementary Fig. 6a-b). Using imaging and flow cytometry, we found that pharmacologic inhibition or knockout of *DGAT1* suppressed lipid droplet biogenesis even when challenged with exogenous fatty acid (Fig. 4a and Supplementary Fig. 6b). Next, we tested if disrupting lipid droplet biogenesis would affect melanoma progression in vivo by knocking out *dgat1a* in our TEAZ model of melanoma (Fig. 4b). Interestingly, knockout of *dgat1a* showed no difference in tumor size at early time points suggesting its loss has minimal effect in tumor initiation. In contrast, loss of *dgat1a* led to significant reduction in tumor area at later time points (Fig. 4c). Overall, this suggests that lipid droplets play a role in later stages of tumor growth and progression.

**Fig. 4:**
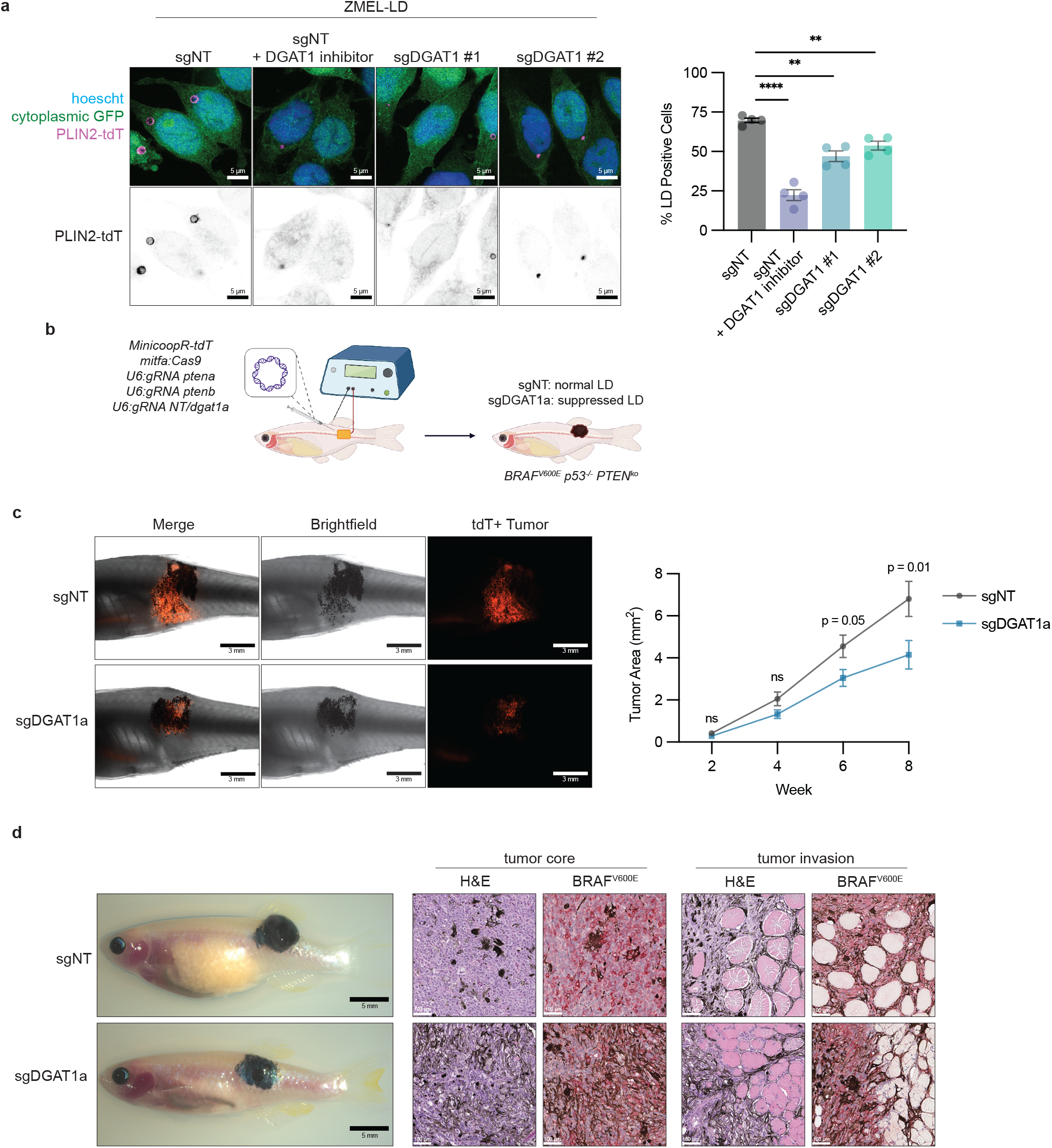
Knockout of DGAT1 suppresses lipid droplet formation and tumor progression. a, Representative fluorescent images of ZMEL-LD lipid droplets marked by PLIN2-tdTOMATO. Cells were incubated with 100 µM oleic acid for 24 hours and lipid droplets were quantified via flow cytometry (mean ± SEM, n = 4 biologically independent experiments). Statistics via two-tailed t-test with Holm-Sidak correction. b, Schematic of zebrafish TEAZ. Transgenic *casper*;*mitfa*:*BRAF^V600E^*;*p53^-/-^*zebrafish were injected with tumor initiating plasmids and electroporated to generate BRAF^V600E^ p53^-/-^ PTEN^ko^ melanomas with normal or suppressed lipid droplet formation. c, Representative images of zebrafish flank with TEAZ generated tumors. Corresponding quantification of tumor area via image analysis as described in Methods (mean ± SEM, n = 3 biologically independent injections, sgNT n = 45, sgDGAT1a n = 49). Statistics via Mann-Whitney U test at each time point. d, Representative whole animal and histological images from week 12 TEAZ generated tumors. Histology shows melanoma morphology at the core of the tumor and at muscle layer tumor invasion. ns p>0.05, ** p<0.01, **** p<0.0001.

Despite the reduction in tumor progression, the *dgat1a* knockout tumors still continued to grow, albeit at a reduced rate. Consistent with this, hematoxylin and eosin (H&E) and BRAF^V600E^ staining showed both control and *dgat1a* knockout tumors could form advanced melanomas capable of tumor invasion beyond hypodermal layers like muscle (Fig. 4d). To gain further insight into the mechanisms sustaining cell growth in *dgat1* deficient tumors, we dissected and sorted tdTomato+ melanoma cells from control and *dgat1a* knockout tumors for RNA-seq (Fig. 5a and Supplementary Data 2). Targeted sequencing of the *dgat1a* locus confirmed the presence of insertions and deletions (indel)^39^ specifically in the *dgat1a* knockout tumors (Supplementary Fig. 6c). Bulk RNA-sequencing of the control versus *dgat1a* knockout melanoma cells showed a significant reduction in *dgat1a* mRNA expression in the knockout tumors (Fig. 5b), as expected. We performed GSEA and observed that the top 5 negatively enriched pathways are cell cycle pathways, consistent with the reduced tumor size in *dgat1a* knockout tumors and reduced proliferative capacity (Fig. 4c and Fig. 5c). However, we also saw upregulation of key de novo cholesterol and fatty acid synthesis genes such as *fasn*, *soat1*, *sqleb*, *hmgcra* suggestive of a compensatory response to dysregulated lipid homeostasis from loss of lipid droplets^40^ (Fig. 5b).

**Fig. 5:**
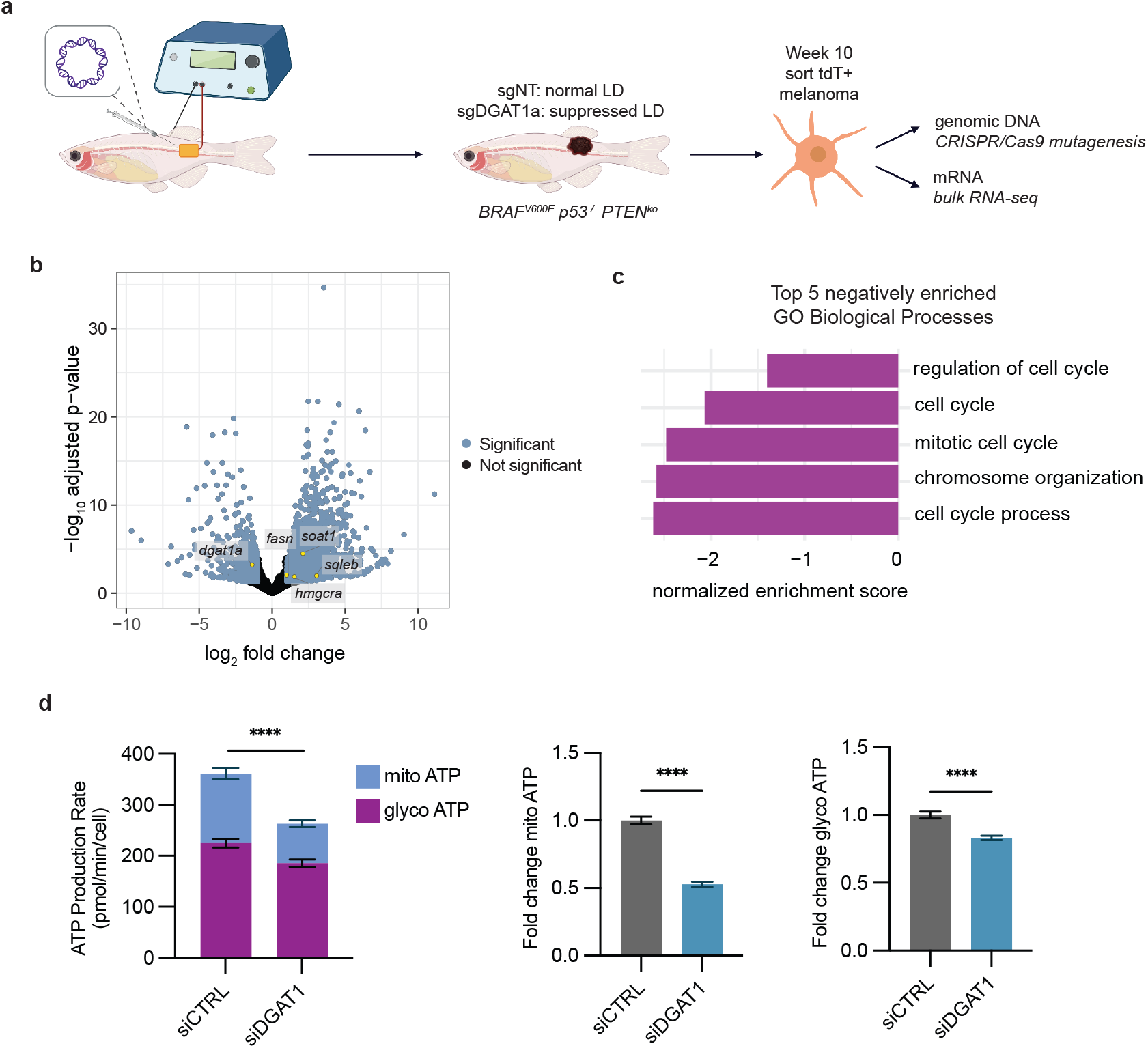
Lipid droplet suppression results in dysregulated metabolic homeostasis. a, Schematic of zebrafish TEAZ and sort for genomic DNA and mRNA. Transgenic *casper*;*mitfa*:*BRAF^V600E^*;*p53^-/-^*zebrafish were injected with tumor initiating plasmids and electroporated to generate *BRAF^V600E^ p53^-/-^ PTEN^ko^* melanomas with normal or suppressed lipid droplet formation. Melanoma cells were sorted to extract genomic DNA and mRNA. b, Volcano plot of bulk RNA-seq showing differential gene expression in sgDGAT1a compared to sgNT melanomas. Blue marks genes significantly upregulated (log_2_ fold change > 1 and Benjamini-Hochberg adjusted p-value<0.05). Yellow marks select fatty acid and cholesterol synthesis genes significantly up- or downregulated. c, Top 5 negatively enriched GO Biological Processes in sgDGAT1a tumors with Benjamini-Hochberg adjusted p-values<0.01. d, Total ATP production rate in human A375 cells divided by contribution from mito ATP (mitochondrial respiration) and glyco ATP (glycolysis). Fold change in mito ATP or glyco ATP quantified by normalizing ATP production rate values to average ATP production rate in siCTRL (mean ± SEM, n = 3 biologically independent experiments). Statistics via two-tailed t-test with Welch correction. **** p<0.0001.

In addition to this compensatory lipid based mechanism, we also considered the possibility that *dgat1a* deficiency leads to glycolytic and oxidative metabolic rewiring. Recent work has shown that lipid droplets are necessary to preserve mitochondrial oxidative function especially during periods of nutrient stress^32,37^. To test whether lipid droplet loss disrupts glycolytic and mitochondrial oxidative function in melanoma cells, we knocked down *DGAT1* in human A375 cells and performed the Seahorse ATP Rate Assay (Supplementary Fig. 6d). We found that knockdown of *DGAT1* was sufficient to reduce total ATP cellular production (Fig. 5d). While *DGAT1* knockdown significantly reduced ATP production from both glycolysis (13% reduction) and mitochondrial respiration (47% reduction), there was a larger effect in mitochondrial ATP production (Fig. 5d). Collectively, these findings indicate that loss of lipid droplets most prominently affects oxidative metabolism, with a smaller effect on glycolytic metabolism.

## Discussion

Phenotypic heterogeneity through non-genetic reprogramming is increasingly recognized as a mechanism for survival in tumors and a therapeutic barrier in melanoma^41^. One example of this reprogramming is metabolic rewiring in which cancer cells metabolically adapt to changing microenvironments and stressors^42^. Metabolic profiling across melanoma cell lines has shown that melanomas have the capacity to simultaneously perform glycolysis and oxidative metabolism even under stressors like hypoxia^43^. However, oxidative metabolism produces significantly more ATP than glycolysis, providing necessary building blocks for cell growth and survival^43^. Previous reports have demonstrated that rewiring to increase oxidative metabolism can drive drug resistance and metastasis in melanoma^44–46^. One mechanism for this increased oxidative metabolic phenotype is mediated by the melanocyte master regulator, MITF, which regulates expression of the metabolic transcriptional coactivator, PGC1ɑ^47^. Conversely, PGC1ɑ can regulate MITF to induce melanogenesis^48^. Our results are consistent with previous studies implicating a direct relationship between melanocytic identity and oxidative metabolism.

Another critical aspect to cancer proliferation is acquisition of substrates to support the energetically expensive process of sustained cellular growth^42^. Many reports support the concept that melanoma cells actively scavenge lipids which can be utilized for processes like membrane formation and energy production^27,28,49,50^. However, lipid accumulation can lead to lipotoxicity. Thus, lipid droplets are critical for storing and facilitating fatty acid release when needed for biosynthetic or energetic purposes^31,32^. Furthermore, lipid droplet biogenesis through DGAT1 has been shown as essential to maintaining mitochondrial health in cancer^37,38^. Here, we show that targeting DGAT1 impairs lipid droplet biogenesis consequently leading to suppressed tumor growth and metabolic dysfunction.

Beyond cellular metabolism, lipid droplet accumulation is tied to cell fate as seen in neural stem and progenitor cells and colorectal cancer stem cells^51,52^. We found that lipid droplets increase with acquisition of more differentiated transcriptional identity in melanoma cells. Altogether, this brings forward the question of how lipid droplets reflect cellular differentiation across different cell types. Recent evidence has shown that lipid droplets maintain physical contacts with the mitochondria, endoplasmic reticulum, lysosome, Golgi apparatus, and peroxisome to function as an antioxidant organelle and mediate inter-organelle transport of macromolecules^30,53^. While our work focused on DGAT1, profiling of the lipid droplet proteome has revealed a surprisingly diverse breadth of metabolic, signaling, trafficking, and membrane organization proteins specifically embedded in the lipid droplet membrane^54^. Future studies will be needed to determine whether perturbing lipid droplet proteins disrupts communication between organelles during cellular demands of tumor progression. Our findings place lipid droplets at the center of oxidative metabolism and lipid regulation, presenting an attractive target at the intersection of metabolic processes necessary for sustained growth in melanoma.

---

**Supplementary Fig. 1:**
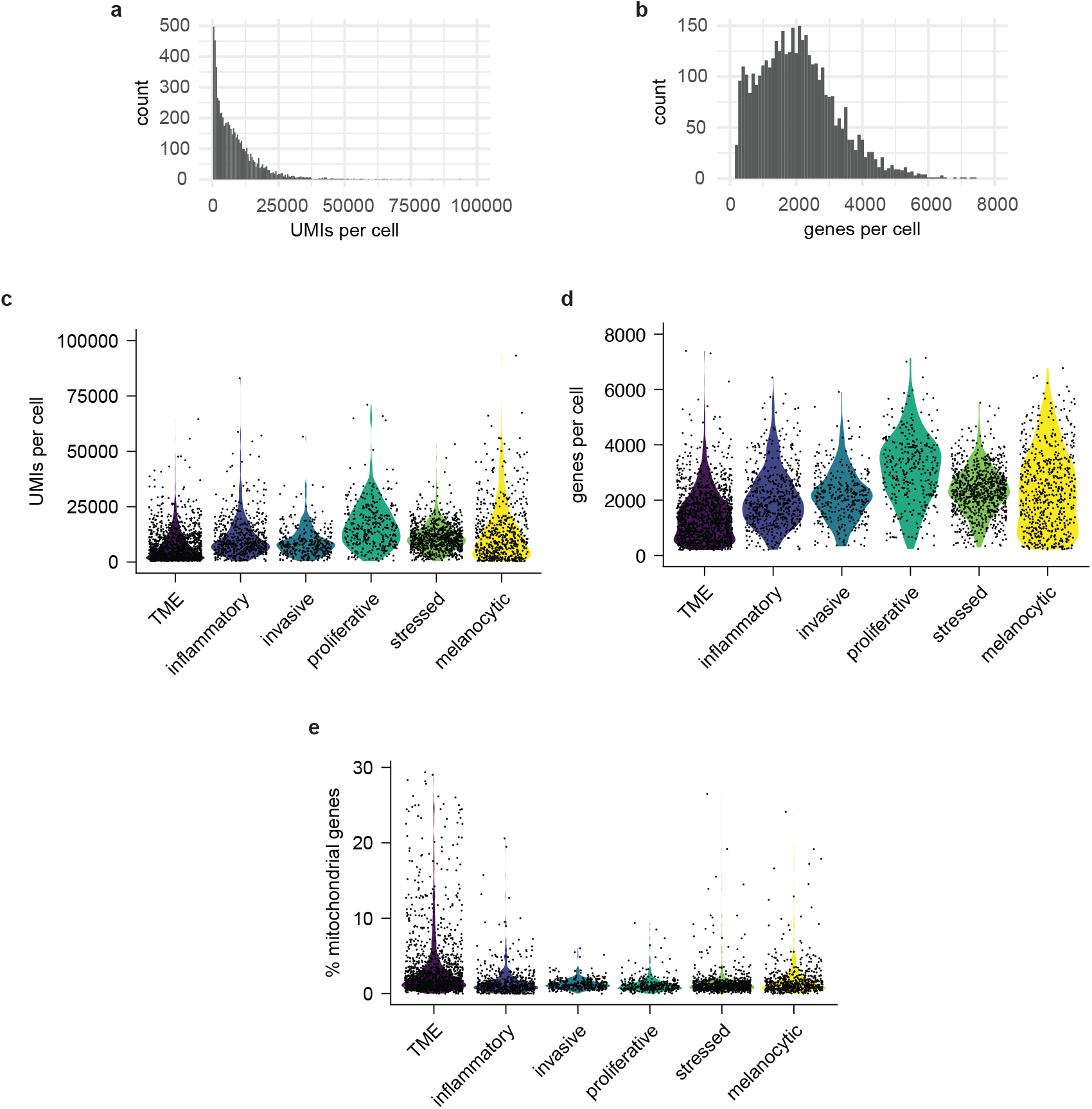
scRNA-seq data statistics. a-b, Histograms of unique molecular identifiers (UMIs): a, and number of unique genes per cell: b, in the total data set of 3968 zebrafish melanoma and TME cells. c-d, Violin plot of UMIs per cell: c, unique genes per cell: d, and percent mitochondrial genes per cell: e, in each cluster assignment.

**Supplementary Fig. 2:**
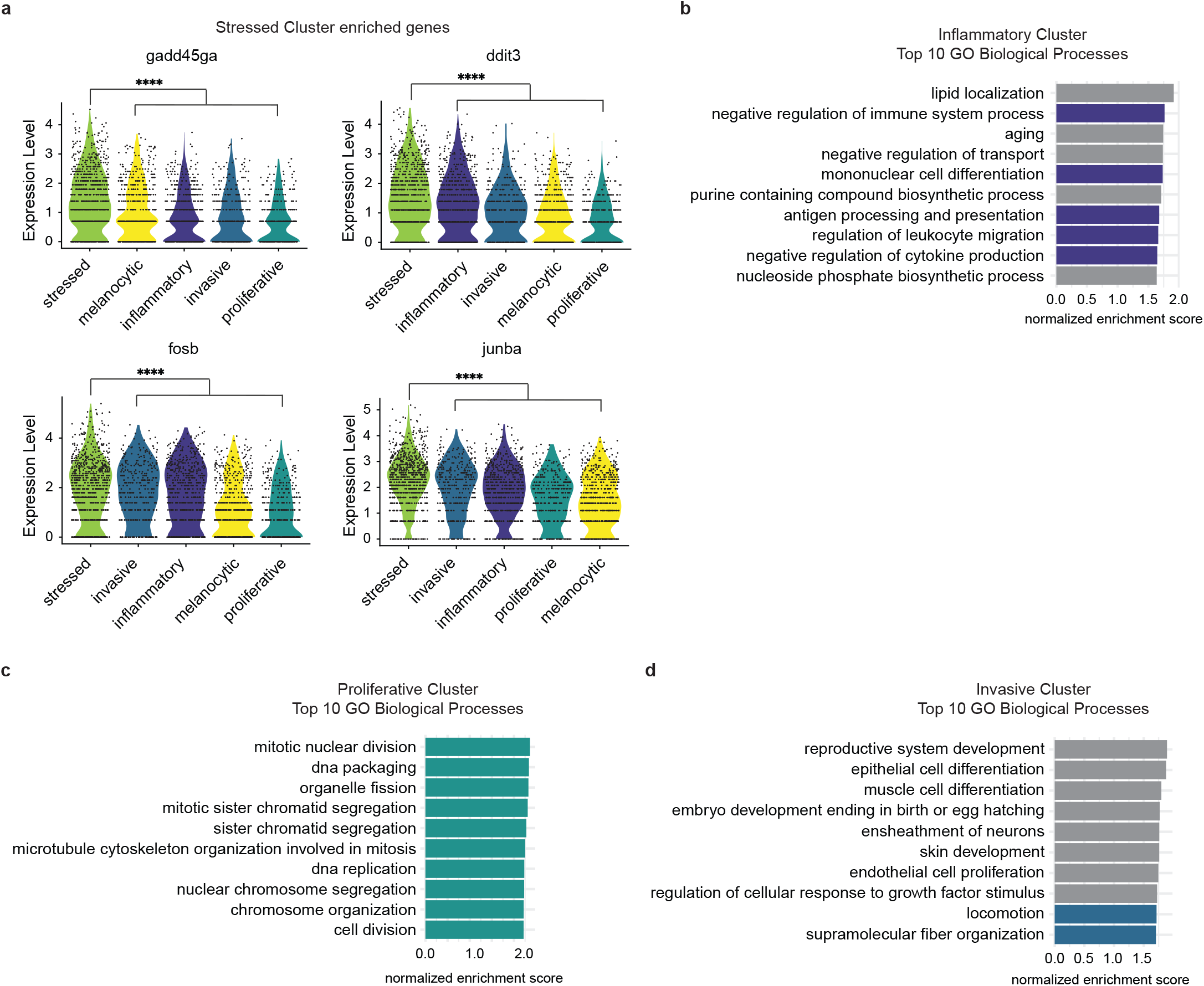
Expression of differentially expressed genes and pathways in stressed, inflammatory, proliferative, and invasive melanoma clusters. a, Violin plot of genes associated with cell stress that are differentially upregulated in the stressed cluster. Statistics via Wilcoxon rank-sum test^22^. **** p<0.0001. b-d, Top 10 enriched GO Biological Processes in inflammatory: b, proliferative: c, and invasive: d, clusters. Pathways are color coded based on relevance to cluster annotation in Fig. 1b.

**Supplementary Fig. 3:**
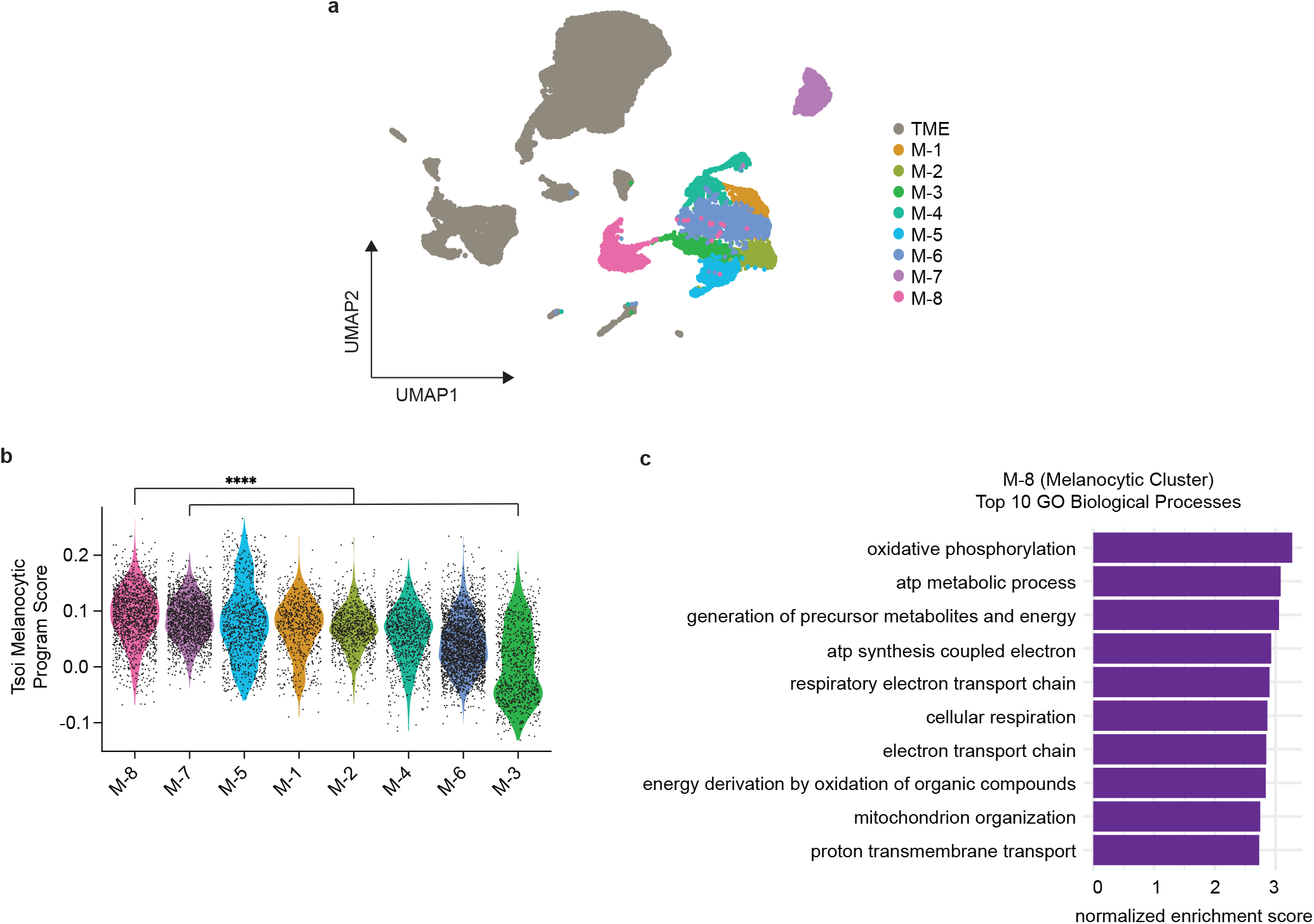
Melanocytic cell state enriches for oxidative metabolic pathways in human melanoma brain and leptomeningeal metastases. a, UMAP projection of human melanoma brain and leptomeningeal metastases from Smalley et al.^24^. Unsupervised clustering^22^ identified eight different melanoma clusters and TME cells as indicated. b, Melanoma cluster 8 (M-8) most enriches for Tsoi Melanocytic Program^4^ module score. Adjusted p-values calculated using Wilcoxon rank-sum test with Holm correction. **** p<0.0001. c, Top 10 enriched GO Biological Processes with Benjamini-Hochberg adjusted p-values<0.01 in M-8 (Melanocytic Cluster) showing oxidative metabolic pathways.

**Supplementary Fig. 4:**
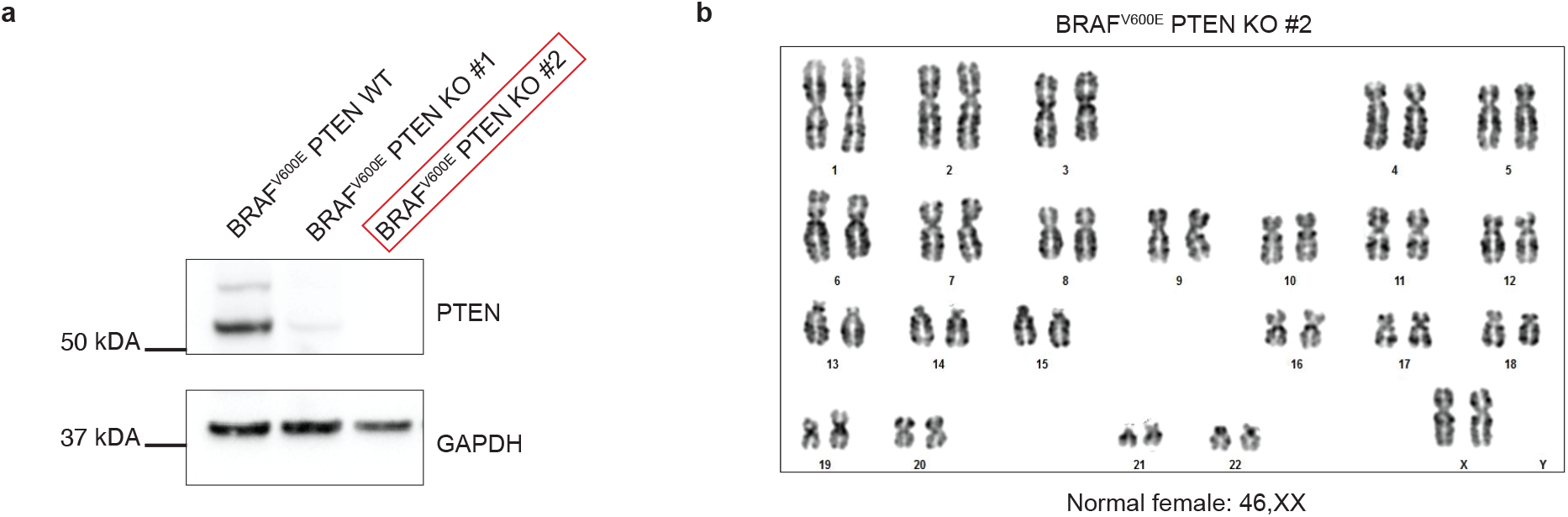
Human pluripotent stem cells (hPSC) PTEN knockout line. a, The doxycycline (dox)-inducible BRAF^V600E^ hPSC line^13^ was genetically engineered to be knockout (KO) for PTEN. Western blotting validates the full loss of PTEN in the KO line #2. b) Karyotypic analysis of the dox-inducible BRAF^V600E^ PTEN KO hPSC line used in this study shows no major chromosomal abnormalities.

**Supplementary Fig. 5:**
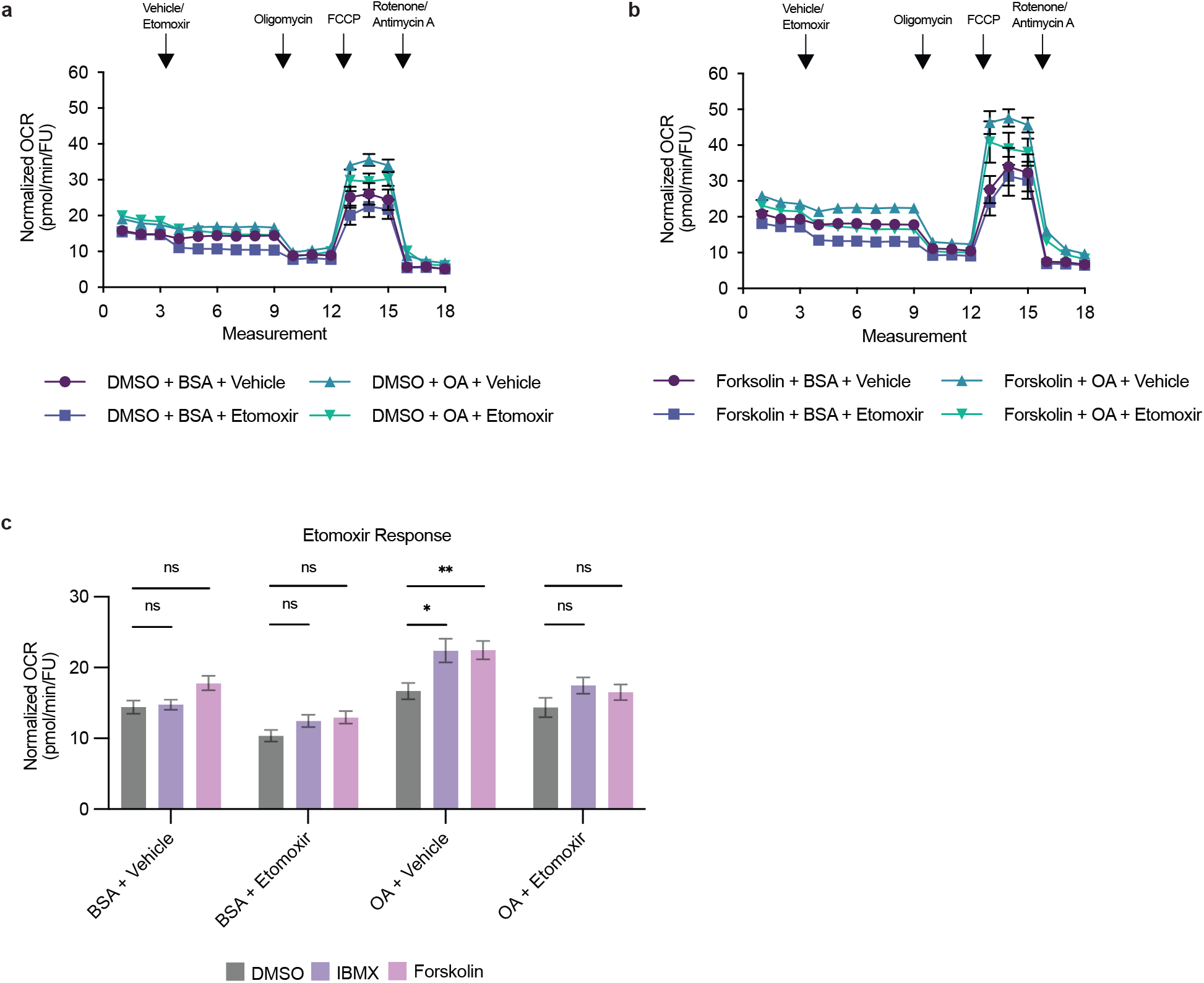
Melanocytic cells utilize fatty acids for beta-oxidation. a-b, Representative normalized OCR measurements for FAO stress test in DMSO (a) or Forskolin (b) treated human A375 cells (mean ± SEM, n = 3 biologically independent experiments). Measurements are plotted separately for ease of visualization. c, Etomoxir response derived from normalized OCR values from measurement 9 in a-b and Fig. 3a in human A375 cells (mean ± SEM, n = 3 biologically independent experiments). Statistics via two-tailed t-test with Holm-Sidak correction. ns p>0.05, * p<0.05, ** p<0.01.

**Supplementary Fig. 6:**
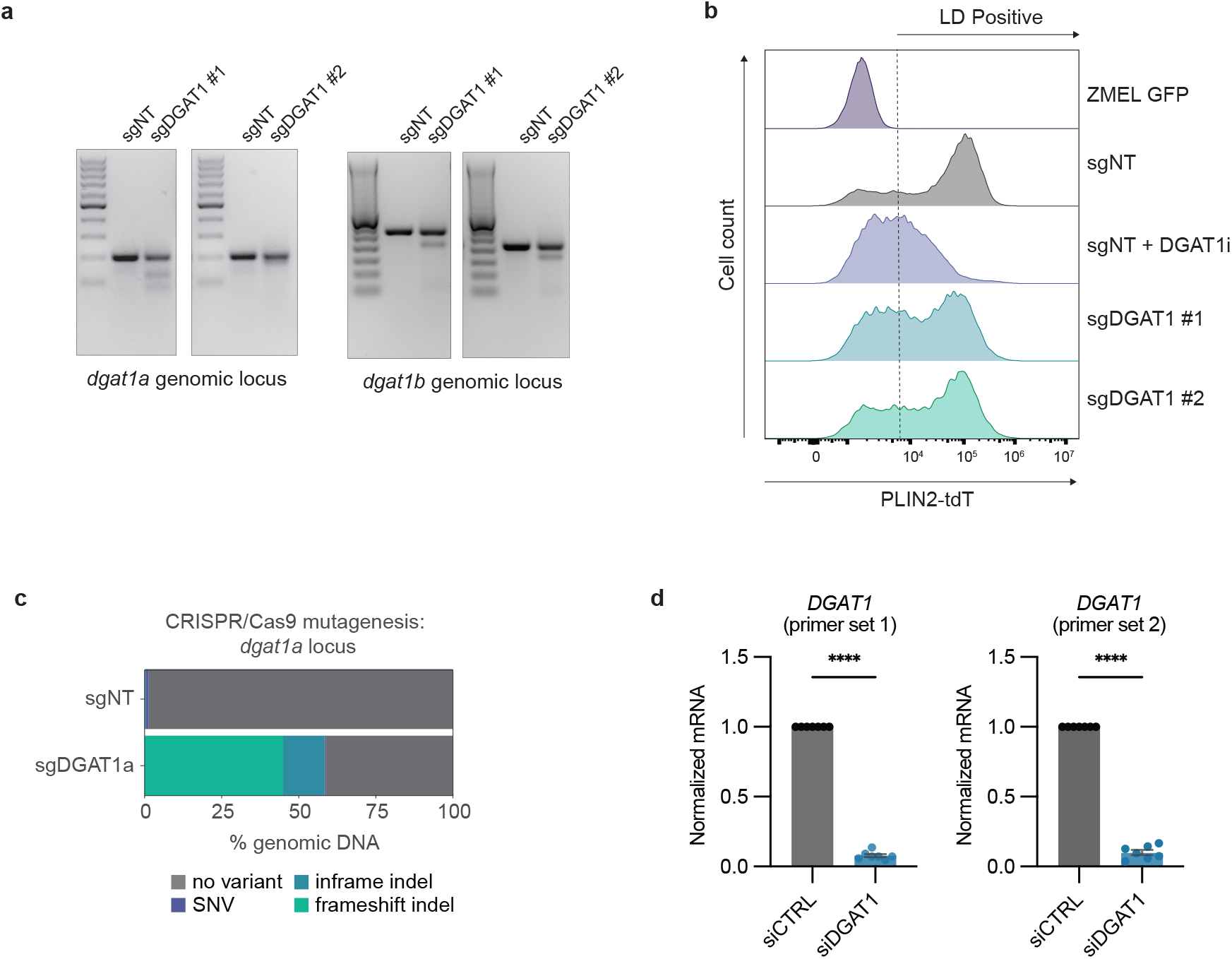
Validation of *dgat1a* and DGAT1 perturbation in zebrafish and human melanoma cells. a, Surveyor nuclease assay for sgDGAT1 guide sets demonstrating indels introduced at the *dgat1a* and *dgat1b* genomic loci. b, Histogram of PLIN2-tdTomato expression in ZMEL-LD cells with DGAT1 perturbation. ZMEL GFP cells devoid of tdTomato expression are negative control for PLIN2-tdTomato. c, Bar plot of CRISPR/Cas9 mutagenesis in *dgat1a* locus of sorted zebrafish melanomas as indicated from Fig. 5a. d, qRT-PCR for *dgat1* in human A375 cells 72 hours post-transfection with control siRNA (siCTRL) or siDGAT1(mean ± SEM, n = 3 biologically independent experiments). Statistics via two-tailed t-test with Welch correction. **** p<0.0001.

## Acknowledgements

We thank the Aquatics Services and Systems facility, A. Afolalu, and M. Shepard for zebrafish maintenance. The Integrated Genomics Organization and Single Cell Research Initiative with R. Chaligne, O. Chaudhary, and N. Sohail provided technical support for bulk and single-cell RNA-sequencing. The Flow Cytometry Core Facility provided technical support for FACS experiments. The Molecular Cytology Core Facility (P30CA008748) provided confocal imaging support. Donald B. and Catherine C. Marron Cancer Metabolism Center, J. Cross, and S.J. Raman provided technical and conceptual support for Seahorse metabolic assays.

## Author contributions

D.L. and R.M.W. developed the experiments and interpreted results. D.L. performed most experiments and collected and analyzed data. D.L., E.M., Y.M. and S.S. performed zebrafish related experiments. Y.M. assisted with computational analyses. C.L. and T.H. assisted with human cell line related experiments. A.B. and L.S. performed and provided hPSC experiments and reagents. D.L and R.M.W. wrote the manuscript. R.M.W. acquired funding for the project. All authors read and edited the manuscript.

## Declarations

R.M.W. is a paid consultant to N-of-One Therapeutics, a subsidiary of Qiagen. R.M.W. receives royalty payments for the use of the casper line from Carolina Biologicals. L.S. is co-founder and consultant of BlueRockTherapeutics and is listed as inventor on a patent application by Memorial Sloan Kettering Cancer Center related to melanocyte differentiation from human pluripotent stem cells (WO2011149762A2). D.L., E.M., A.B., Y.M., C.L., T.H. and S.S. declare no competing interests.

## Methods

### Zebrafish husbandry

Zebrafish were housed in a temperature- (28.5°C) and light-controlled (14 hr on, 10 hr off) room. Zebrafish were anesthetized using Tricaine (MS-222) with a stock of 4 g/l (protected for light) and diluted until the fish was immobilized. All procedures were approved by and adhered to Institutional Animal Care and Use Committee (IACUC) protocol #12-05-008 through Memorial Sloan Kettering Cancer Center.

### Cell culture

Human melanoma A375 cells were obtained from ATCC and routinely confirmed to be free from mycoplasma. Cells were maintained in a 37°C and 5% CO^2^ humidified incubator. Cells were maintained in DMEM (Gibco, 11965) supplemented with 10% FBS (Gemini Bio, 100-500) and 1x penicillin/streptomycin/glutamine (Gibco, 10378016). Cells were used for experiments until passage 25.

Zebrafish ZMEL-LD cells were generated as recently described^33^ and routinely confirmed to be free from mycoplasma. Cells were maintained in a 28°C and 5% CO^2^ humidified incubator. Cells were maintained in DMEM supplemented with 10% FBS, 1x penicillin/streptomycin/glutamine, and 1x GlutaMAX (Gibco, 35050061). Cells were used for experiments until passage 25.

### Zebrafish melanoma generated by TEAZ

Melanomas were generated by Transgene Electroporation in Adult Zebrafish as previously described^13,21^. To generate BRAF^V600E^ p53^-/-^ PTEN ^ko^ melanomas, adult transgenic zebrafish (*casper (mitfa^-/-^;mpv17^-/-^)*;*mitfa*:*BRAF^V600E^*;*p53^-/-^*) were injected with the following tumor initiating plasmids: MinicoopR-tdT (250 ng/µL), *mitfa:Cas9* (250 ng/µL), *U6:sgptena* (23 ng/µL), *U6:sgptenb* (23 ng/µL) and Tol2 (57 ng/µL). For *dgat1a* knockout experiments, zebrafish were injected with an additional guide plasmid of *U6:sgNT* (23 ng/µL) or *U6:sgdgat1a* (23 ng/µL). Adult male and female zebrafish were anesthetized in tricaine and injected with 1 µL of tumor initiating plasmids below the dorsal fin, electroporated using the CM 830 Electro Square Porator from BTX Harvard Apparatus, and recovered in fresh water. For *dgat1a* knockout experiments, zebrafish were imaged every two weeks using brightfield and fluorescence imaging using a Zeiss AxioZoom V16 fluorescence microscope. To quantify tumor area, images were analyzed in MATLAB R2020a by quantifying pixels positive for melanin and tdTomato.

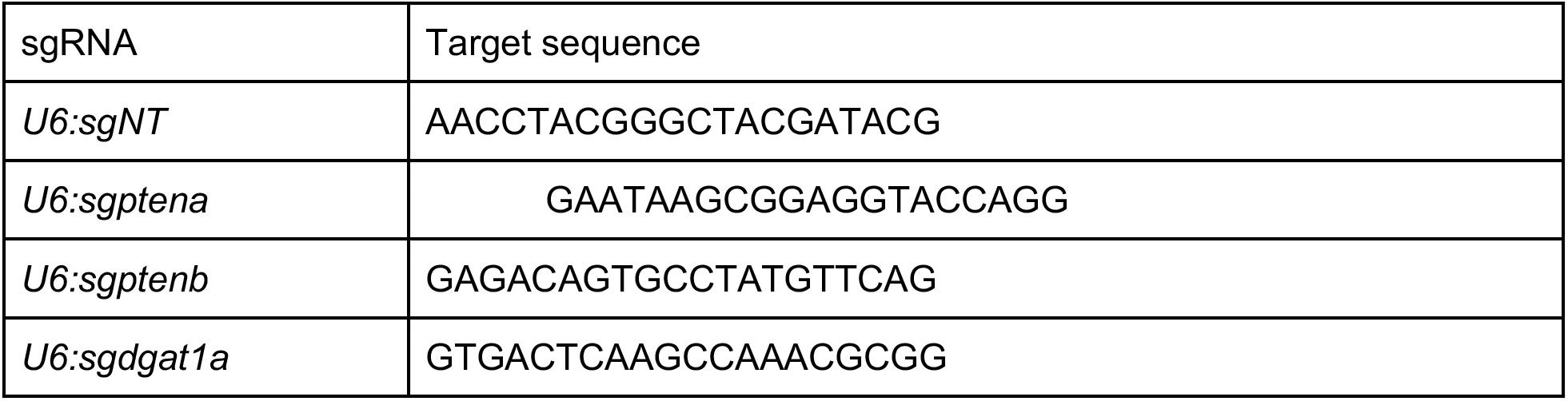

### Zebrafish histology and immunohistochemistry

Zebrafish were sacrificed in an ice bath for at least 15 min. Zebrafish were fixed in 4% paraformaldehyde for 72 hr at 4°C, washed in 70% ethanol for 24 hours, and then paraffin embedded. Fish were sectioned at 5 mm and placed on Apex Adhesive slides, baked at 60°C, and then stained with hematoxylin & eosin or antibodies against BRAF^V600E^ (1:100, Abcam, ab228461) using the BOND Polymer Refine Red Detection Kit (Leica Biosystems, DS9390). Whole slide scanning was performed on an Aperio AT2 digital whole slide scanner (Leica Biosystems). Histology was performed by Histowiz.

### scRNA-sequencing of zebrafish melanomas

Six zebrafish (3 male and 3 female) with melanoma (3 months post-TEAZ) were anesthetized and sacrificed using Tricaine. Tumor and adjacent tissue was dissected and dissociated using 0.16 mg/mL Liberase (Sigma-Aldrich, #5401020001) in 1x PBS and gentle pipetting with a wide-bore P1000 for 30 minutes at room temperature. Dissociation was terminated with addition of 250 µL FBS and samples were filtered through a 70 µm filter. Male and female zebrafish were labeled by sex using the 3’ CellPlex Kit Set A (10x Genomics, 1000261) per the manufacturer’s instructions. Resulting cell pellets were resuspended in 1x PBS and 1% UltraPure BSA (Thermo-Fisher, AM2616) and passed through a 40 µm filter into 5 ml polystyrene tubes for FACS. Cells were sorted to remove debris and doublets using BD FACSAria III cell sorter.

Library preparation and sequencing were done by the Single Cell Research Initiative and Integrated Genomics Organization at MSKCC. For cell encapsulation and library preparation, droplet-based scRNA-seq was performed on approximately 5900 cells using the Chromium Single Cell 3’ Library and Gel Bead Kit v3 and Chromium Single Cell 3’ Chip G (10x Genomics) into a single v3 reaction. GEM generation and library preparation were performed according to manufacturer instructions. Libraries were sequenced on a NovaSeq6000. Sequencing parameters: Read1 28 cycles, i5 10 cycles, i7 10 cycles, Read2 90 cycles. Sequencing depth was approximately 51,000 reads per cell.Sequencing data was aligned to our reference zebrafish genome using CellRanger 6.0.2 (10x Genomics)^55^.

Data was processed using R version 4.0.5 and Seurat version 4.0.2^22^. Cells with fewer than 200 unique genes and mitochondrial genes above 30% were filtered. Expression data was normalized using SCTransform with principal component analysis and UMAP dimensionality reduction performed at default parameters. Clustering was performed using the Seurat function FindClusters with resolution of 0.4. Cluster annotation for zebrafish cell-type specific marker genes as done previously using FindAllMarkers^8,55^.

Differentially expressed genes for pathway analysis was performed using the Seurat function FindMarkers comparing the tumor clusters. Ribosomal genes and genes with p-values<0.05 were filtered out. Ortholog mapping between zebrafish and human was performed with DIOPT^13,55^. Gene set enrichment analysis was performed using fgsea 1.16.0 using the MSigDB GO biological processes (GO.db 3.12.1)^55^. Calculation of Tsoi Melanocytic Program^4^ Score was determined using the Seurat AddModuleScore function with default parameters.

### Re-analysis of Smalley et al.^24^ human melanoma scRNA-seq

Data was processed using R version 4.0.5 and Seurat version 4.0.2^22^. Counts matrix was obtained from GEO (GSE1744401) and Seurat object was created with default parameters as described previously^55^. Calculation of Tsoi Melanocytic Program^4^ Score was determined using the Seurat AddModuleScore function with default parameters.

### Bulk RNA-sequencing of zebrafish melanomas

Zebrafish tumors were dissected and sorted for tdTomato+ cells as described above. mRNA was extracted and prepared with SMARTer Universal Low Input RNA Kit for Sequencing (Takara) for 100 bp paired-end sequencing on the NovaSeq 6000 by the Integrated Genomics Organization at MSKCC. Sequencing reads underwent quality control with FASTQC 0.11.9, trimming with TRIMMOMATIC 14.0.1 and aligned using Salmon 1.4.0 to the *danio rerio* GRCz11.

Data analysis was conducted in R version 4.0.5. Differential expression was calculated using DESeq2 1.30.1 using default parameters. Significant differentially expressed genes were called if log_2_ fold change>1 and adjusted p-value<0.05. Ortholog mapping between zebrafish and human was performed with DIOPT^13,55^. Gene set enrichment analysis was performed using fgsea 1.16.0 using the MSigDB GO biological processes (GO.db 3.12.1)^55^.

### Generation of PTEN knockout line

sgRNAs were cloned into the PX458 Cas9-GFP vector and introduced into dox-inducible BRAF^V600E^ hPSC by nucleofection as previously described^13^. Cells were FACS sorted 24 hours post nucleofection, and individually seeded on a mouse embryonic fibroblast feeder layer in the presence of 10 mM ROCK-inhibitor in knockout serum replacement stem cell media^56^ for two weeks. ROCK-inhibitor was removed from culture media after 4 days. Clones were transferred to vitronectin-coated plates and further maintained in E8. Full loss of PTEN expression was finally validated by Western blotting (anti-PTEN antibody, Cell Signaling, 9559S, 1:1000). sgRNAs come from Cederquist et al.^57^:

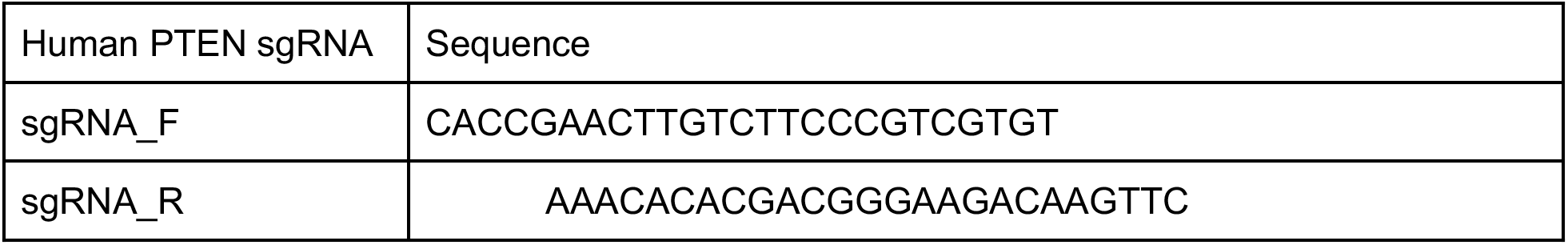

### Melanoblast differentiation protocol

Dual SMAD inhibition protocol was performed as previously described^11^ and melanocytes differentiation was executed previously^13^.

In brief, the dox-inducible BRAF^V600E^ PTEN KO hPSC were plated as a high-density monolayer with 150,000 cells per cm2. This is a lower density than that one used for the WT cells, because of the higher proliferation rates due to the PTEN knockout. It is important to induce BRAF^V600E^ only at the end of the differentiation. This otherwise impairs hPSC differentiation.

<u>Day -1</u>: Plate hPSCs on Matrigel in E8 medium with 10μM ROCKi (R&D, 1254).

<u>Day 0-2</u>: Change media every day with E6 media containing 1ng/ml BMP4 + 10μM SB + 600nM CHIR.

<u>Day 2-4</u>: Change media with E6 media containing 10μM SB + 1.5μM CHIR.

<u>Day 4-6</u>: Change media with E6 media containing 1.5μM CHIR.

<u>Day 6-11</u>: Change media every day with E6 media containing 1.5μM CHIR + 5ng/ml BMP4 100nM EDN3.

### Flow cytometry associated cell sorting

Dox-inducible BRAF^V600E^ PTEN KO hPSC-derived melanoblasts were sorted at day 11 of differentiation using a BD-FACS Aria6 cell sorter at the Flow Cytometry Core Facility of MSKCC. The cells in differentiation were initially dissociated into single cells using Accutase (Innovative Cell Technologies, 397) for 20 minutes at 37°C and then stained with a conjugated antibody against cKIT (Anti-Hu CD117 (cKIT) (APC), Invitrogen #17-1179-42) and P75 (anti-CD271 (FITC), BioLegend #345104). Cells double positive for FITC (P75) and APC (cKIT) were sorted and 4, 6-diamidino-2-phenylindole (DAPI) was used to exclude dead cells.

### Melanoblasts expansion

At day 11, melanoblasts were aggregated into 3D spheroids (2 million cells/well) in ultra-low attachment 6-well culture plates (corning, 3471). Cells were expanded for maximum 7 days and then used for the Seahorse experiments.

#### Melanoblasts media

Neurobasal media (gibco, 21103-049)

1mM L glutamine (gibco, 25030-081)

0.1 mM MEM NEAA (gibco, 11140-050)

FGF2 10ng/ml (R&D, 233-FB/CF)

CHIR 3uM (R&D, 4423)

B27 supplement (gibco, 12587-010)

N2 supplement (gibco, 17502-048)

### Melanocyte differentiation

Upon FACS sorting, P75^+^cKIT^+^ melanoblasts were plated onto dried PO/Lam/FN dishes. Cells were fed with melanocyte medium every 2 to 3 days. Cells were passaged once a week at a ratio of 1:6, using accutase for 20 min at 37°C for cell detachment. Mature melanocytes at day 100 were used for the seahorse experiments.

#### Melanocyte media (∼1L)

Neurobasal media 500ml (gibco, 21103-049)

DMEM/F12 500ml (gibco, 11330-032)

SCF 25ng/ml (R&D, 255-SC-MTO)

cAMP 250uM (Sigma, D0627)

FGF2 5ng/ml (R&D, 233-FB/CF)

CHIR 1.5uM (R&D, 4423)

BMP4 12.5ng/ml (R&D, 314-BP)

EDN3 50nM (Bachem, 4095915.1000)

25 ml FBS (R&D, S11150H)

2.5 ml penicillin/streptomycin (gibco, 15140-122)

2 ml L-Glutamine (gibco, 25030-081)

B27 supplement (gibco, 12587-010)

N2 supplement (gibco, 17502-048)

### Induction of melanocytic cell state in A375 cell line

Human A375 cells were trypsinized, centrifuged at 300g for 3 minutes, and counted for viability then 500,000 viable cells per well were seeded in a 6 well TC treated plate. After 6 hours and confirming cell attachment, media was aspirated and fresh media was added with either DMSO, 200 µM IBMX (Cayman Chemical, 13347), or 20 µM Forskolin (EMD Millipore, 344270). Cells were incubated in drugs for 72 hours then harvested for qRT-PCR or Seahorse Mito Stress Test.

### qRT-PCR

Total RNA was isolated using the Quick-RNA Miniprep Kit (Zymo, R1055) according to the manufacturer instructions. cDNA was synthesized using SuperScript IV First-Strand Synthesis System (Thermo Fisher, 18091200) and qPCR was performed using Applied Biosystems PowerUp (Thermo Fisher, A25742). Results were normalized to the *beta-actin* housekeeping gene.

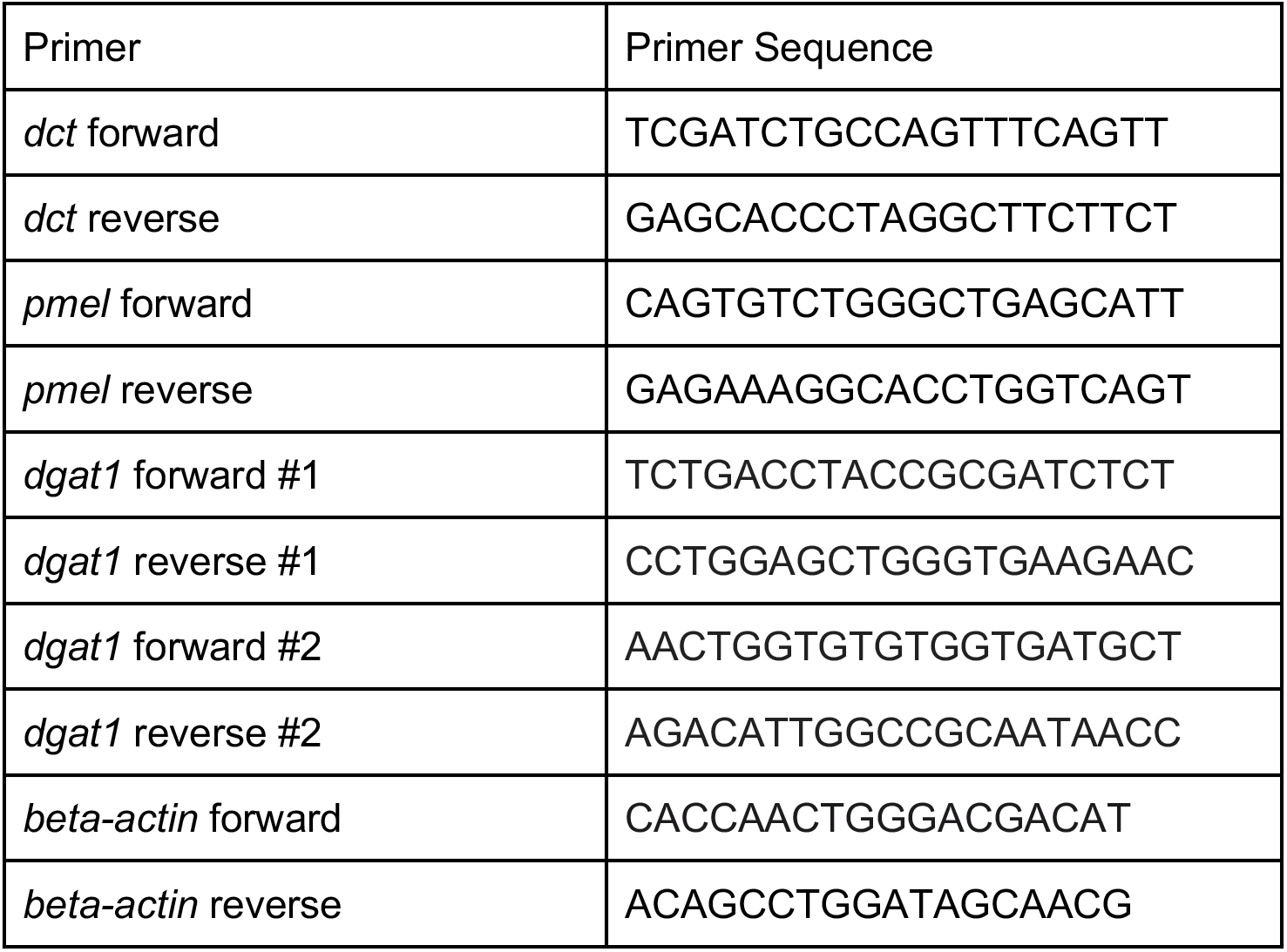

### Seahorse Mito Stress Test

The Seahorse XF Mito Stress the Seahorse XFe96 Analyzer. For melanoblasts and melanocytes, cells 30,000 melanoblasts and 10,000 melanocytes were seeded per well in a XF cell plate as previously described^13^. For human A375 cells, cells were treated with DMSO, IBMX, or Forskolin as described above then cells were trypsinized and resuspended in drug containing media at 30,000 cells per well in a XF cell plate coated with 0.05% poly-L-lysine then incubated overnight (Sigma-Aldrich, 4707).

Cells were incubated in XF Mito Stress Test assay medium (Seahorse XF DMEM medium, pH 7.4, 10 mM glucose, 2 mM glutamine, 1 mM sodium pyruvate) for 1 hour prior to measurement in a CO^2^ free incubator at 37°C. During assay run, cells were exposed to 2.0 µM oligomycin, 2.0 µM FCCP and 0.5 µM rotenone/antimycin A. OCR and ECAR were normalized to nuclei fluorescence unit (FU) via SYTO 24 (Thermo Fisher, S7559) or protein via Pierce BCA Protein Assay Kit (Thermo Fisher, 23227) as indicated. Experimental measurements were analyzed using the Agilent Wave software.

### Fatty acid uptake

A375 cells were seeded at 30,000 cells per well in a 96 well TC treated black microplate (Greiner Bio-One, 655090) and incubated with DMSO, 200 µM Forskolin or 20 µM IBMX for 24 hours. Cells were washed 3 times with DMEM 1% FBS and incubated in DMEM 1% FBS with DMSO, Forksolin or IBMX for one hour. Lipid uptake was measured using the QBT Fatty Acid Uptake Assay (VWR, 10048-826) using the BioTek Synergy plate reader. Fluorescence at 485 nm excitation and 528 nm emission was measured every 50 seconds for 30 minutes. Fold change fatty acid uptake was measured by normalizing fluorescence at each time point to the fluorescence at the start of the assay.

### Seahorse Fatty Acid Oxidation Test

A375 cells were trypsinized, centrifuged at 300g for 3 minutes, and counted for viability then 500,000 viable cells per well were seeded in a 6 well TC treated plate. After 6 hours and confirming cell attachment, media was aspirated and fresh media was added with either DMSO, 200 µM IBMX, or 20 µM Forskolin. Cells were incubated in drugs for 48 hours and then tested for fatty acid oxidation in a protocol adapted from the Seahorse XF Palmitate Oxidation Stress Test. Cells were trypsinized and resuspended in DMEM 10% FBS at 30,000 cells per well in a XF cell plate coated with 0.05% poly-L-lysine (Sigma-Aldrich, 4707). Cells were allowed to adhere to the plate for 5 hours then washed twice with Seahorse XF DMEM medium, pH 7.4. Cells were incubated overnight in DMSO, 200 µM IBMX, or 20 µM Forskolin in nutrient limited media: Seahorse XF DMEM medium, pH 7.4, 5 mM glucose, 1 mM glutamine, 1% FBS, and 0.5 mM carnitine (Fisher Scientific, AC230280050). In addition, cells were supplemented with either 150 µM BSA (Sigma-Aldrich, A1595) or 150 µM oleic acid conjugated to BSA (Sigma-Aldrich, O3008).

We used the Seahorse XF Long Chain Fatty Acid Oxidation Stress Test kit (Agilent, 102720-100). Cells were incubated in assay media supplemented with 150 µM BSA or oleic acid conjugated to BSA as fatty acid substrate in the following formulation: Seahorse XF DMEM medium, pH 7.4, 5 mM glucose, 1 mM glutamine, 0.5 mM carnitine. Cells were incubated for 1 hour prior to measurement in a CO^2^ free incubator at 37°C. During assay run, cells were exposed to 10 µM etomoxir, 1.5 µM oligomycin, 2.0 µM FCCP and 0.5 µM rotenone/antimycin A. OCR and ECAR were normalized to nuclei fluorescence unit (FU) via SYTO 24 (Thermo Fisher, S7559). Experimental measurements were analyzed using the Agilent Wave software.

### siRNA knockdown

Dharmacon siGENOME siRNA reagents were used according to manufacturer instructions. 200,000 A375 melanoma cells were seeded in antibiotic free DMEM 10% FBS in a 6 well TC treated plate with 5 µM siGENOME Non-targeting siRNA SMARTPool (Horizon Discovery, D-001206-13-05) or siGENOME DGAT1 siRNA SMARTPool (Horizon Discovery, M-003922-02-0005) with DharmaFECT1 Transfection Reagent (Horizon Discovery, T-2001-01). Media was replaced with regular media including antibiotics 24 hours post-transfection. After 48 hours, cells were harvested for Seahorse ATP Rate Assay or total RNA extraction.

### Seahorse ATP Rate Assay

The Seahorse XF Real-Time ATP Rate Assay (Agilent, 103592) was performed using the Seahorse XFe96 Analyzer. Cells were trypsinized and seeded at 30,000 cells per well in a XF cell plate coated with 0.05% poly-L-lysine then incubated overnight. Cells were incubated in assay medium (Seahorse XF DMEM medium, pH 7.4, 10 mM glucose, 2 mM glutamine, 1 mM sodium pyruvate) for 1 hour prior to measurement in a CO^2^ free incubator at 37°C. During assay run, cells were exposed to 1.5 µM oligomycin and 0.5 µM rotenone/antimycin A. OCR, ECAR, and Proton Efflux Rate (PER) were normalized to nuclei fluorescence unit (FU) via SYTO 24 (Thermo Fisher, S7559). Experimental measurements were analyzed using the Agilent Wave software generating the mito ATP and glyco ATP measurements.

### Lipid droplet staining and imaging

For human A375 cells, 40,000 cells were seeded in each well of the Millicell EZ slide 4-well (EMD Millipore, PEZGS0416) with regular media and DMSO, IBMX or Forskolin. After 72 hours, cells were fixed with 4% formaldehyde for 15 minutes, washed with 1x PBS and permeabilized in 0.1% Triton-X 100 in PBS for 30 minutes. Cells were washed, blocked with 10% goat serum (Thermo Fisher, 50-062Z), and incubated overnight at 4°C in 1:100 dilution of rabbit anti-PLIN2 (Proteintech, 15294-1-AP). Cells were washed with incubate for 1 hour at room temperature in 1:500 goat anti-rabbit Alexa Fluor 555 (Cell Signaling Technology, 44135), 1:400 Alexa Fluor 488 Phalloidin (Thermo Fisher, A12379) and 1:2000 Hoechst 33342 (Thermo Fisher, H3570).

For zebrafish ZMEL-LD cells, 250,000 cells were seeded in each well of the Millicell EZ slide 4-well and incubated with 100 µM of oleic acid for 24 hours. For cells treated with DGAT1 inhibitor, cells were treated with 20 µM T-863 (Cayman Chemicals, 25807). Cells were fixed with 4% formaldehyde for 15 minutes, washed with 1x PBS and stained with 1:2000 Hoescht for 10 minutes.

Slides were mounted in Vectashield Antifade Mounting Media (Vector Laboratories, H-1000). Cells were imaged on the Zeiss LSM 880 inverted confocal microscope with AiryScan using a 63x oil immersion objective. Confocal stacks were visualized in FIJI 2.1.0 and A375 melanoma lipid droplets were counted using FIJI.

### CRISPR/Cas9 knockout in ZMEL-LD

The Alt-R CRISPR/Cas9 System (Integrated DNA Technologies) and Neon Transfection System were used according to manufacturer protocols for CRISPR/Cas9 knockout in ZMEL-LD cells. Knockout was validated by harvesting genomic DNA with the DNeasy Blood & Tissue Kit (Qiagen, 69506), *dgat1a* and *dgat1b* genomic loci were PCR amplified with Platinum SuperFi II PCR Master Mix (Thermo Fisher, 12368010) and indels detected using the Surveyor Mutation Kit (Integrated DNA Technologies, 706020). Cells were incubated overnight with 100 µM oleic acid then analyzed for lipid droplets via flow cytometry as previously described^33^. Briefly, cells were stained for viability with 1:1000 DAPI, data acquired using the Beckman Coulter CytoFLEX Flow Cytometer (Beckman Coulter) and analyzed using FlowJo 10.8.1 (BD Biosciences).

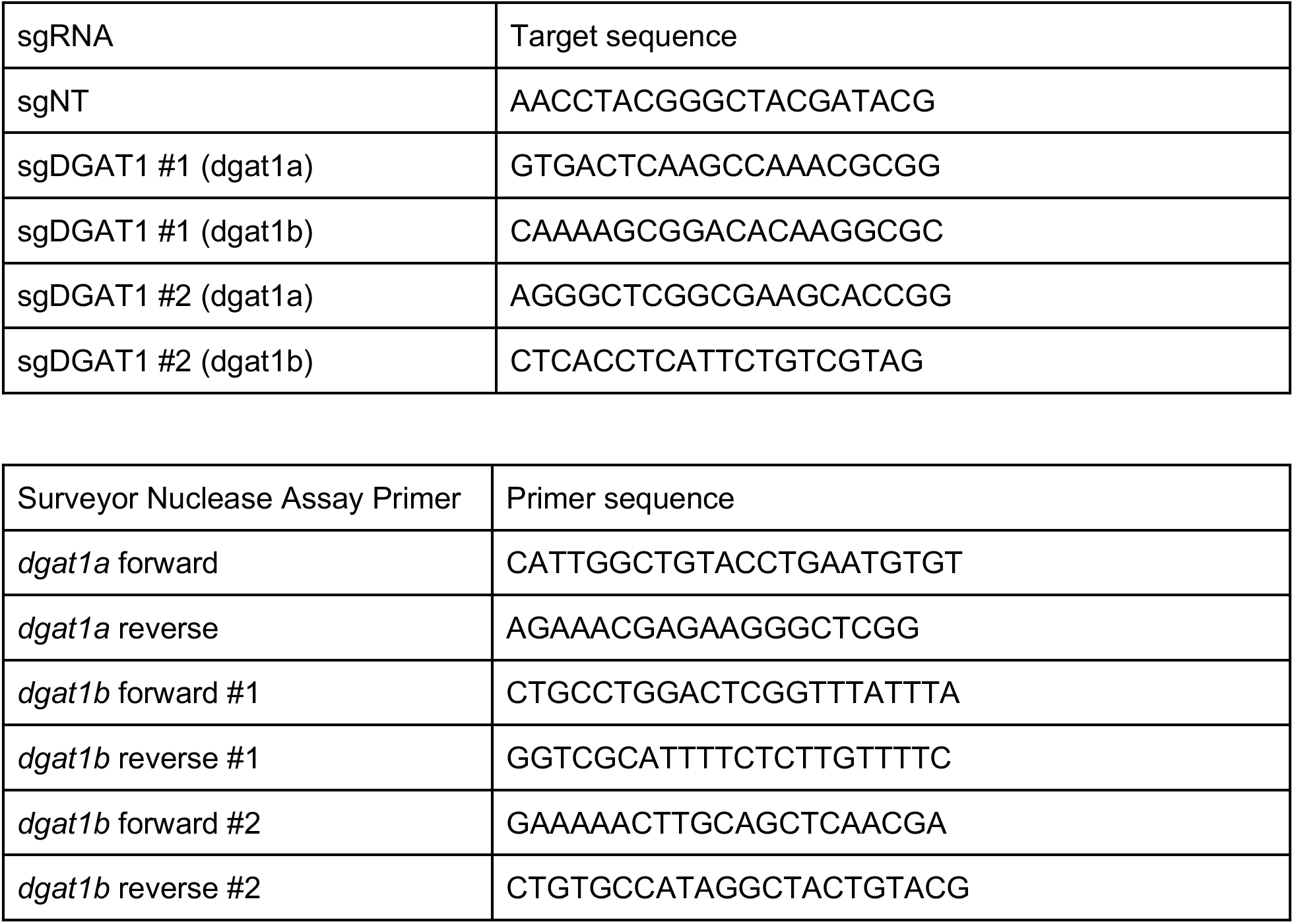

### CrispRVariants knockout

Zebrafish tumors were dissected and sorted for tdTomato+ cells as described above then harvested for genomic DNA with the DNeasy Blood & Tissue Kit and the *dgat1a* locus was PCR amplified using primers from surveyor nuclease assay. DNA was purified using the NucleoSpin Gel & PCR Clean-up Kit (Takara, 740986.20) and 100 bp paired end reads sequenced using Illumina NovaSeq at the Integrated Genomics Operation at MSKCC. Sequences were aligned and percent of indels at the target site was quantified using R version 4.0.5 and CrispRVariants 1.18.0 as previously described^39^.

### Statistics and reproducibility

Statistical analysis and figures were generated in GraphPad Prism 9.1.1, R Studio 4.0.5, and Biorender.com. Image processing and analysis was performed in MATLAB R2020a and FIJI 2.1.0. Statistical tests and p-values are reported in the figure legend for each experiment. Experiments were performed at least three independent times unless noted in the figure legend for each experiment. All sequencing data will be uploaded to the Gene Expression Omnibus database. All other relevant data supporting the key findings of this study are available within the article and its Supplementary Data files or from the corresponding author upon reasonable request.

